# Unlocking HDR-mediated Nucleotide Editing by identifying high-efficiency target sites using machine learning

**DOI:** 10.1101/464610

**Authors:** Aidan R. O’Brien, Laurence O.W. Wilson, Gaetan Burgio, Denis C. Bauer

## Abstract

Editing individual nucleotides is a crucial component for validating genomic disease association. It currently is hampered by CRISPR-Cas-mediated “base editing” being limited to certain nucleotide changes, and only achievable within a small window around CRISPR-Cas target sites. The more versatile alternative, HDR (homology directed repair), has a 4-fold lower efficiency with known optimization factors being largely immutable in experiments. Here, we investigated the variable efficiency-governing factors on a novel mouse dataset using machine learning. We found the sequence composition of the repair template (ssODN) to be a governing factor, where different regions of the ssODN have variable influence, which reflects the underlying biophysical mechanism. Our model improves HDR efficiency by 83% compared to traditionally chosen targets. Using our findings, we develop CUNE (Computational Universal Nucleotide Editor), which enables users to identify and design the optimal targeting strategy using traditional base editing or – for-the-first-time – HDR-mediated nucleotide changes.

CUNE can be run via the web at: https://gt-scan.net/cune

## Introduction

The targeted aproach for disease treatment, precision medicine (1), has greatly benefited from advancements in sequencing technologies, which enable the economical identification of potentially disease causing genetic variants for cancer, neurological and immunological disorders (2–4). However, the function of many of these detected rare variants are unknown, and with current functional assays being costly and time-consuming, precision medicine is bottlenecked by this step. The CRISPR-Cas (Clustered Regularly Interspaced Short Palindromic Repeats) system, a recently developed versatile gene-editing technology, allows researchers to modify DNA or RNA of living cells more easily and hence promises to accelerate knowledge-gain from functional studies.

One of the most efficient methods for modifying single nucleotides using the CRISPR-Cas system is “base editing”. Base editing requires a catalytically inactive Cas9 or a Cas9 nickase mutant. When fused to a cytidine (C) deaminase or adenosine (A) deaminase enzyme, this can convert C•G to T•A, or A•T to G•C, respectively (5,6). Different groups have recently demonstrated CRISPR base editing to achieve efficiencies of 44%–100% (median=82%) in mice, rabbits, rats and human embryos (7–10). The efficiency depends on factors including the surrounding sequence composition, and the position of the point mutation relative to the protospacer adjacent motif (PAM) at the target site. Base editing is only efficient within a tight parameter space, e.g. only ~5 bases at each CRISPR-Cas binding site can be targeted with high efficiency (11). While work is underway to extend the “editing window” (12), there are drawbacks as larger editing windows can result in proximal off-targets, that is, besides the intended target, all or a random selection of nucleotides of the same letter are changed. This together with the restricted range of changes (C → T, G → A, A → G, T → C) limits the application of base editing substantially.

A more-versatile approach to inducing point mutations is via the homology directed repair (HDR) pathway. The HDR pathway is one of two main DNA repair pathways present in organisms from prokaryotes to eukaryotes. HDR accurately repairs Cas9-induced double strand breaks (DSBs) using a homologous DNA template (13,14). Including a mutation in a synthetic DNA template enables HDR to introduce said mutation into the target.

However, inducing point mutations using HDR can be inefficient as HDR is cell-cycle dependent (15) and in direct competition with the error-prone non-homologous end joining (NHEJ) pathway (14,16). While the binding and cutting efficiency of CRISPR-Cas can be predicted with computational tools (17–20), factors influencing the repair pathway choice and hence the potential for successfully introducing a point mutation remain unknown.

Here, we identified factors that influence Cas9-mediated HDR efficiency using machine learning on a novel fit-for-purpose dataset. From these insights, we built a computational tool that allows researchers to introduce a wider range of mutations than base editing, by harnessing the versatility of HDR. CUNE, Computational Universal Nucleotide Editor, is available as a webservice at https://gt-scan.net/cune, giving the user the choice of finding target sites for HDR and base editing.

## Results

### A dataset of genome-wide HDR efficiencies

We curated data from a series of experiments aiming to change a specific nucleotide in mice using Cas9-mediated HDR. To achieve this, we used the CRISPR-Cas9 system to induce a DNA double-strand break (DSB) at a chosen position near the intended point mutation. We used a single-stranded oligodeoxynucleotide sequence (ssODN) template (a point mutation flanked by homology arms, homologous to the PAM strand of the target region) to define the mutation. Table 1 lists statistics on the 30 unique experiments we used to calculate the HDR and NHEJ efficiency.

**Table 1.**
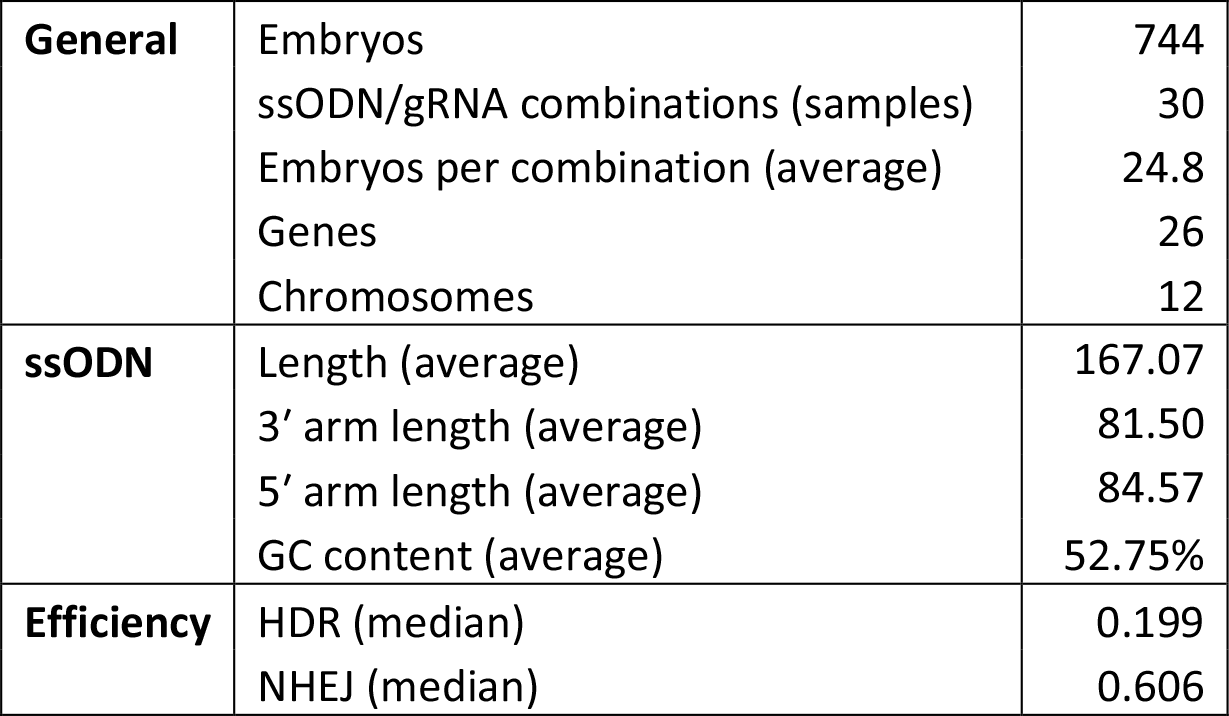
Our dataset includes results from HDR experiments in 744 embryos. This includes 30 ssODN/gRNA combinations (with approximately 25 embryos each). The median HDR and NHEJ efficiency for each ssODN/gRNA combination is 0.199 and 0.606, respectively. The ssODNs have an average length of 167 (with the arm length between 81 and 85 on average).

|  |  |  |
| --- | --- | --- |
| **General** | Embryos | 744 |
|  | ssODN/gRNA combinations (samples) | 30 |
|  | Embryos per combination (average) | 24.8 |
|  | Genes | 26 |
|  | Chromosomes | 12 |
| **ssODN** | Length (average) | 167.07 |
|  | 3' arm length (average) | 81.50 |
|  | 5' arm length (average) | 84.57 |
|  | GC content (average) | 52.75% |
| **Efficiency** | HDR (median) | 0.199 |
|  | NHEJ (median) | 0.606 |

To infer the repair mechanism used, we classified mice carrying the desired mutation as having been repaired via HDR, and mice carrying arbitrary mutations as having been repaired by the error-prone NHEJ. To quantify the HDR efficiency of each target, we divided the number of mice with HDR repairs by the number of mice with any repair (HDR or NHEJ) for each of our 30 targets.

The dataset is a general representation of HDR targets across the mouse genome, with 26 genes targeted across 12 chromosomes. The 30 loci (unique HDR targets) are derived from 126 experiments, with an average of 5.9 mice per locus, i.e. 744 mice (Table 1).

The distribution of efficiencies is shown in Figure 1, clearly demonstrating the 3-fold lower efficiency for HDR (median = 0.199) compared to NHEJ (median = 0.606) when choosing targets traditionally. These values are comparable to previous work (21–23).

**Figure 1.**
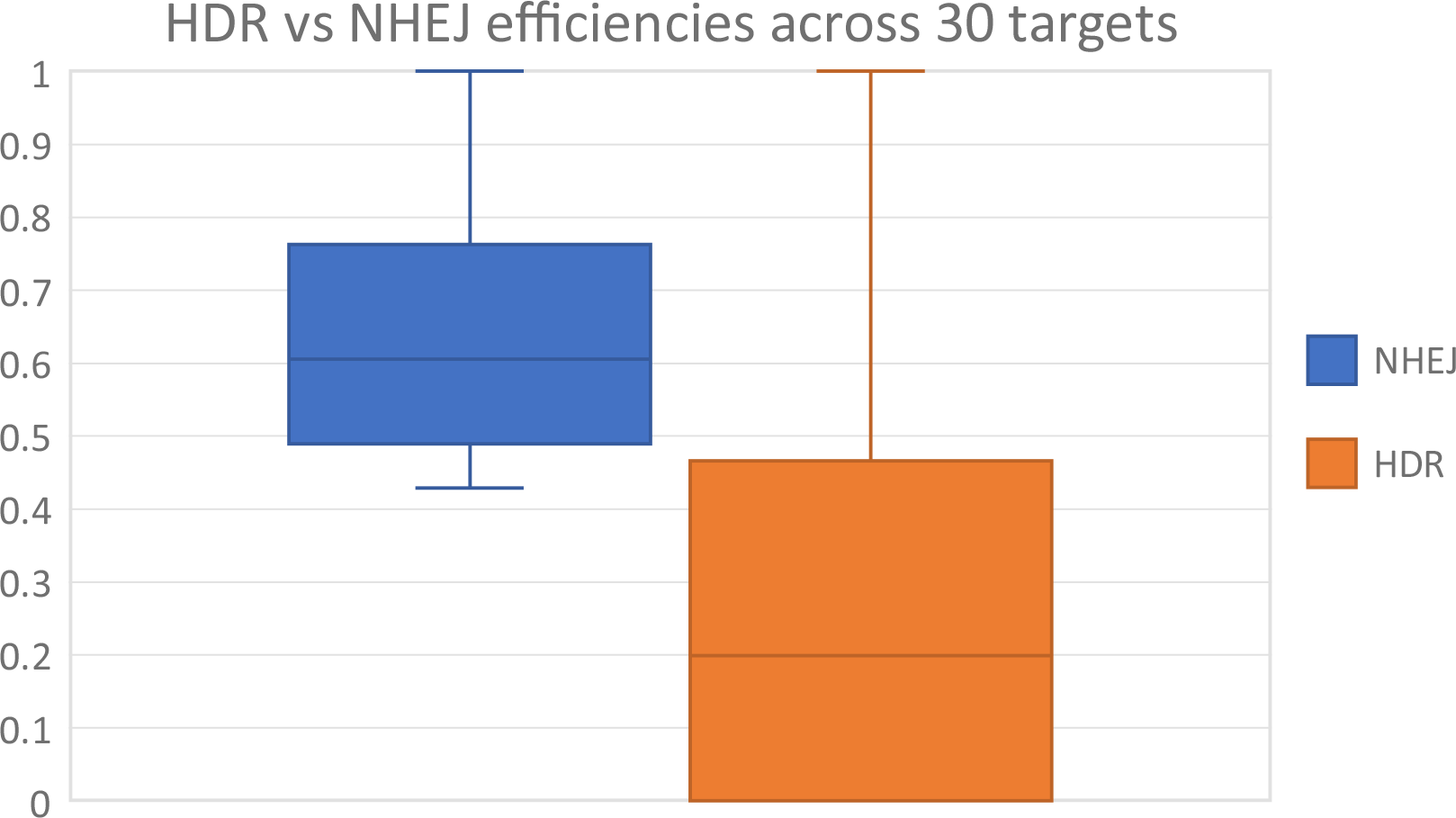
Distributions of HDR and NHEJ efficiencies across the 30 loci. The median HDR efficiency (0.199) is 3-folds lower than the median NHEJ efficiency (0.606). Although HDR (0.466) has a higher interquartile range than NHEJ (0.334), Q3 of HDR is less than Q1 of NHEJ, indicating the inefficiency of HDR versus NHEJ.

To generate the training dataset for the efficiency predictor, we split targets into “high” and “low” HDR efficiency on the median (0.199). This generates a balanced dataset (50% high and 50% low), reducing the potential for classification bias.

### Guide nucleotide composition informs Cas9-mediated HDR insertion rate

Using our dataset, we trained machine learning models to predict CRISPR-Cas9-mediated HDR efficiency for new target sites. We took inspiration from computational methods for predicting generic CRISPR-Cas9 activity, which make predominant use of the nucleotide composition of the gRNA (24). While HDR requires control over the repair-pathway, it may still be predominantly driven by Cas9 activity. We therefore investigated whether the nucleotide composition of the guide is sufficient for predicting knock-in efficiency through HDR.

We used the random forest machine learning algorithm to model the data, and the following results are the average across five cross-validated folds. We trained our first model using the guide nucleotide and dinucleotide composition (G1), and subsequent models on other sequence-based features such as repeats or the type of nucleotides present (G2-G4). The nucleotide composition is the count of each nucleotide (i.e. Ts, Gs, etc.) in the sequence, and the dinucleotide composition is the count of adjacent nucleotides (i.e. TTs, GTs, etc.). We recorded metrics including the out-of-bag (OOB) error, area under the ROC curve, precision, recall and number of samples classified correctly. Out of these four models, the simplest model (G1) of the (di)nucleotide compositions produced the lowest error (OOB = 0.25), where including the pyrimidine/purine composition (G4) resulted in the poorest model (OOB = 0.33) (Table 2).

**Table 2.**
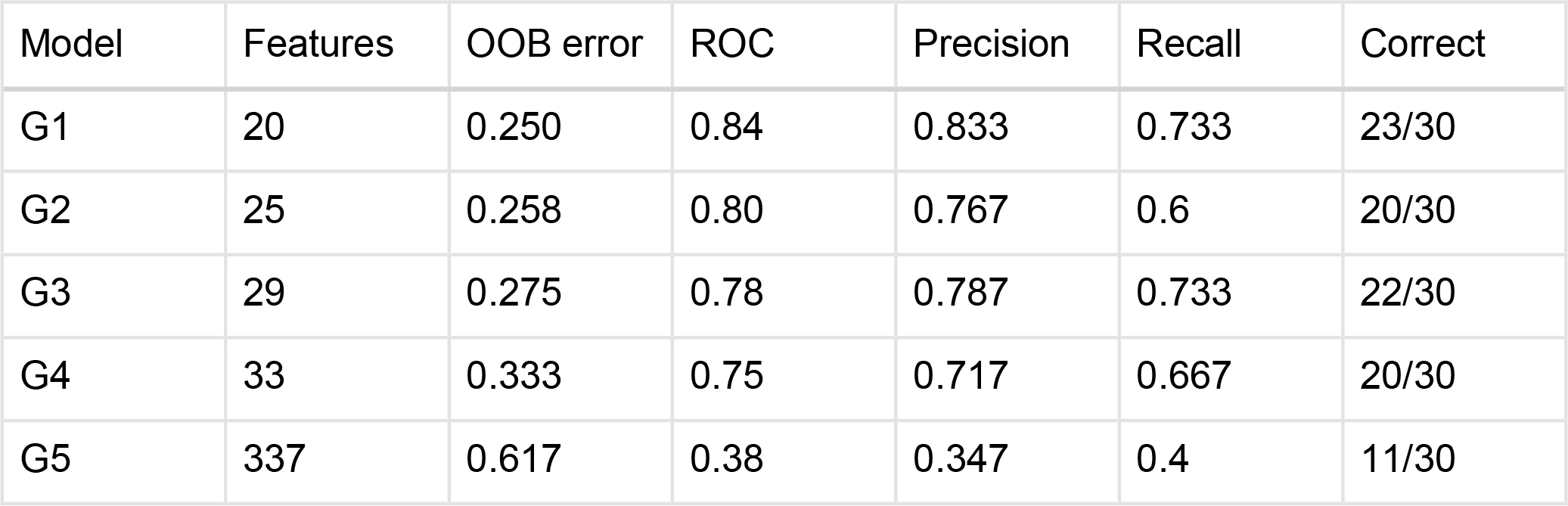
G1: guide nucleotide composition, G2: G1 + generic repeats, G3: G2 + guide AT/CG composition. G4: G3 + guide pyrimidine/purine composition, G5: local guide nucleotides. Metrics are averaged over the five cross-validated folds.

We also built a model on positional nucleotide information (G5). In contrast to G1, which models the position-independent nucleotide count (i.e. how many Cs are in the sequence), G5 models the presence of each nucleotide at each position (i.e. is there a C present at position 1, position 2, etc.). This position-dependent model performs the poorest of all five, with an OOB of 0.617. The poor performance is likely due to the large number of features (337), relative to the sample size of 30. An increase in this ratio generally correlates with the sparsity of the dataset, i.e. the number of zeros. This phenomenon is known as the “curse of dimensionality” and makes it increasingly difficult for machine learning algorithms to find signal in the data (25).

### Mutation-to-cut distance does not improve model accuracy

With evidence that the distance between the cleavage-site and the mutation has an inverse relationship to HDR efficiency (26,27), we retrained the above models with distance as an input feature. We observed the expected inverse relationship between distance and HDR efficiency (Figure 2, fitted quadratic model peaks at the cut site), albeit with a low coefficient of determination (R² = 0.0926). However, we did not observe an improvement in prediction accuracy of the trained model.

**Figure 2.**
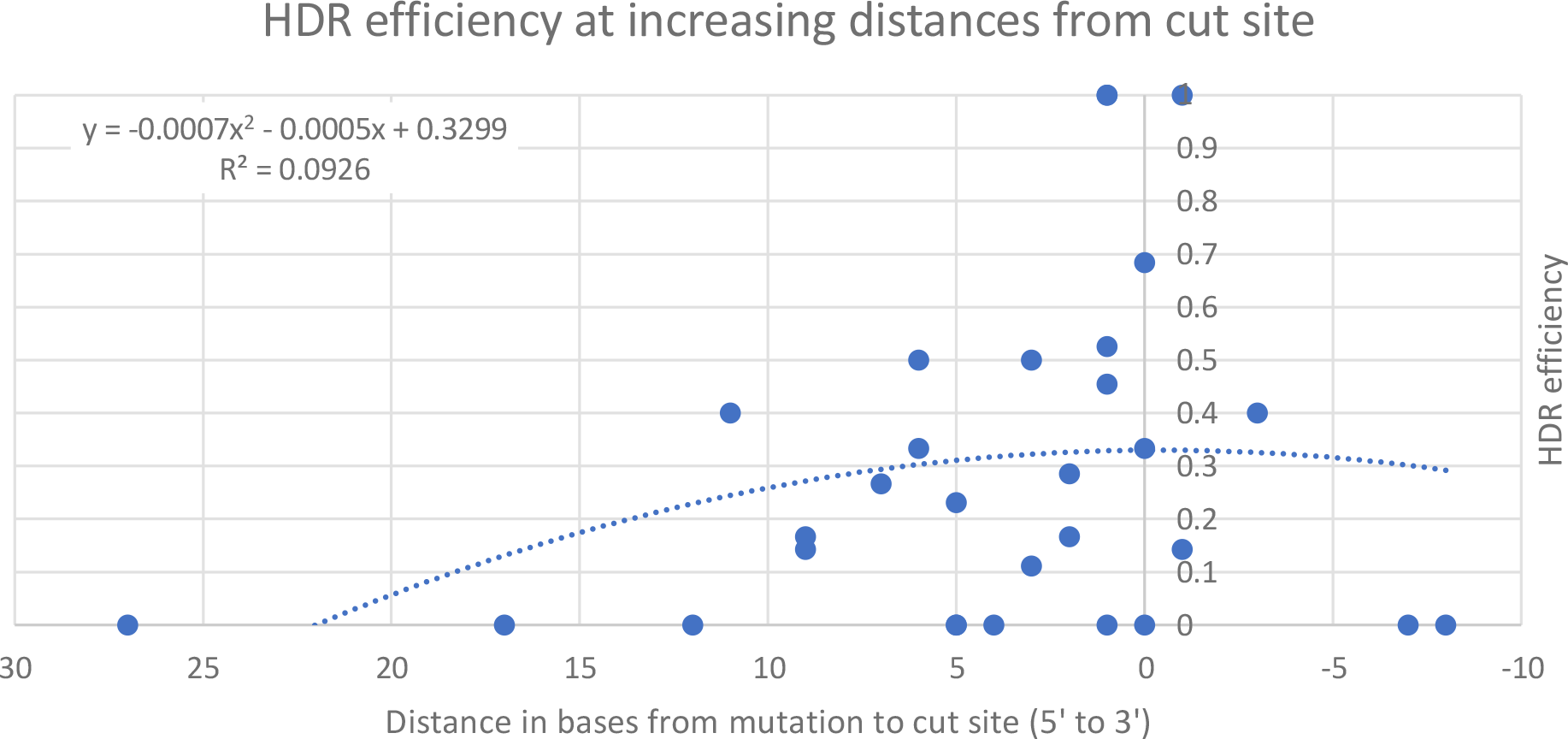
HDR efficiencies for targets plotted against the distance from cut site. Positive values are on the 5′ side of the cut on the PAM strand (5′ – 3′, left to right)

The dataset was designed to capture a wide range of factors, such as gene locus or genomic location influencing HDR efficiency (28). Relative to those other factors, editing distance seems to be a weak modulator of efficiency as editing may inherently fail at inefficient loci, regardless of distance.

### The 3′ homology arm informs HDR activity

Although we demonstrated that the guide sequence influences Cas9-mediated HDR efficiency, it is likely not influencing HDR directly. The observed higher HDR activity is likely an indirect result of higher DSB frequency. We therefore investigated whether modelling more-direct properties of HDR can improve on our previous best model (G1).

Due to the key role of the ssODN in inducing the desired point mutation, we hypothesised that a model trained on the ssODN would be able to accurately differentiate between high- and low-efficiency targets. Models trained on the ssODN capture a larger number of nucleotides than the gRNA models (~167 vs 23), see Figure 3a. However, it appears that this extra information does not contribute to the prediction power, as the ssODN model (O1) performs poorly, with an OOB error of 0.6 (vs. 0.25).

**Figure 3.**
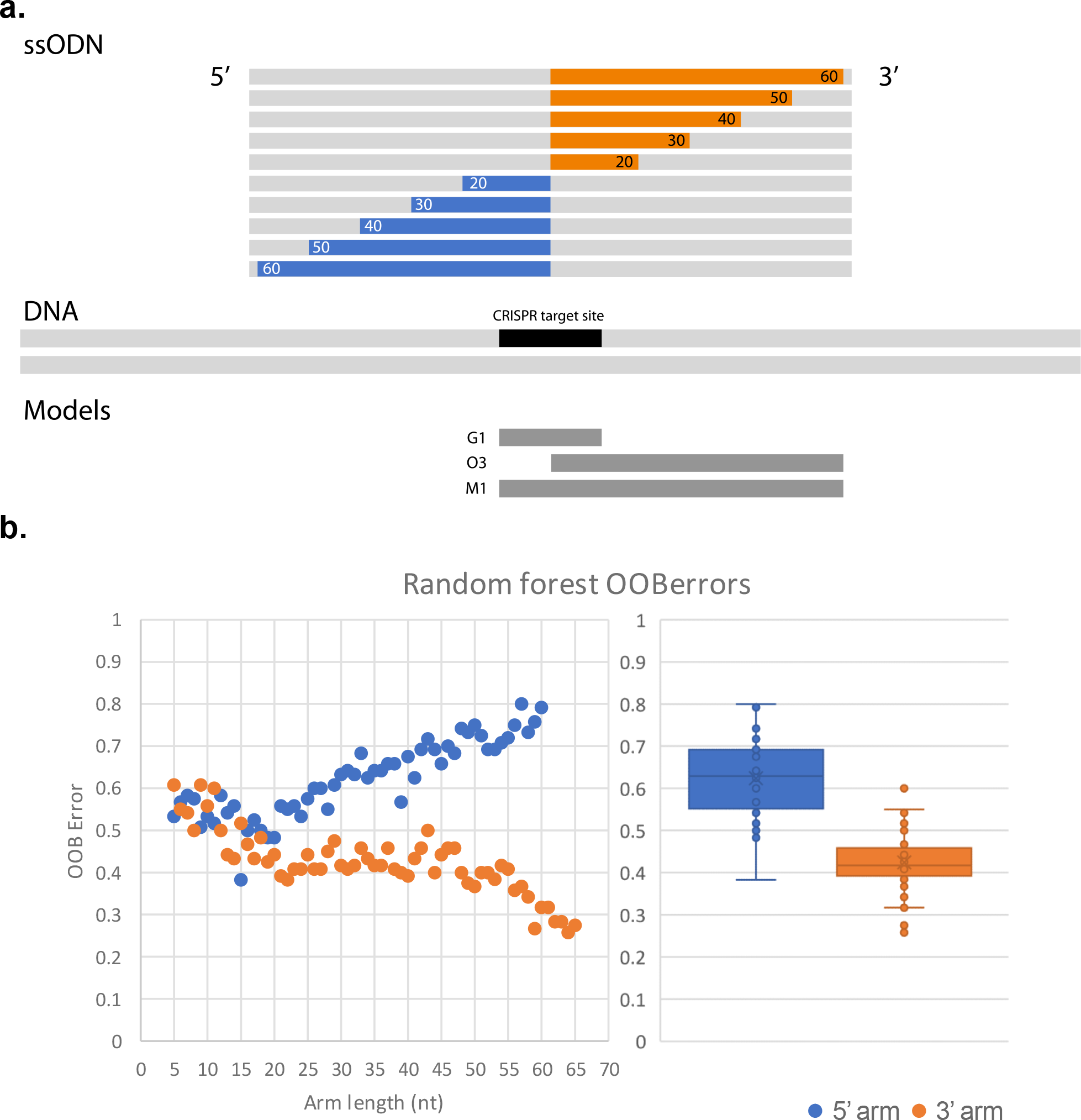
(a) The upper gray bars represent an ssODN. The blue (5’) and orange (3’) bars represent the regions of the ssODN we include in each model. We model regions in increments of 1, but only factors of 10 are displayed for clarity. Each region begins at the center of the ssODN and extends outwards. The lower bar represents the approximate alignment between the ssODN and the CRISPR target. (b) The OOB error for random forest models built on the above regions of the ssODN. Regions start from the center and extend outward. Each point represents the error for a model built on one of these regions. The presented maximum lengths, 60nt for the 5′ arm and 65nt for the 3′ arm, are dictated by the smallest respective arm in the dataset.

Other groups have investigated the influence of the symmetry and length of ssODN arms on HDR activity, drawing the conclusion that asymmetric ssODNs can improve HDR efficiency (29,30). The consensus is that a shorter 3′ arm with a longer 5′ arm is the optimal design for efficient HDR, with one theory being that the shorter 3′ arm allows the ssODN to anneal to the DNA target without requiring further processing or strand invasion (30). Expanding on this, we investigated whether the correlation between HDR efficiency and the ssODN nucleotide composition differs for each arm. We investigated the information content for each arm separately by training different models on each arm. As presented in Table 3, the 3′ arm (homologous to the PAM/non-target strand) informs HDR efficiency, with an OOB of 0.275, while the 5′ arm performs poorly, with an OOB of 0.792.

**Table 3.**
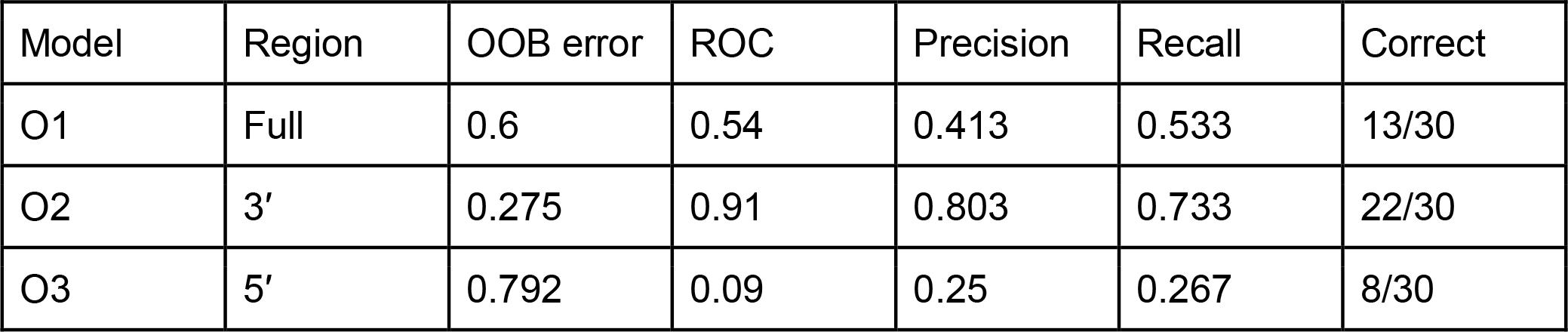
Metrics from three random forest models trained on the nucleotide composition of the ssODN. O1 is trained the full ssODN. O2 is trained on the 3′ arm, and O3 is trained on the 5′ arm (all homologous to the PAM strand).

With a clear difference in predictive power between the two regions of the ssODN (O2 vs O3), we investigated this difference further by modelling regions of each of the arms. Each region starts at the middle of the ssODN, next to the point mutation, and extends outward toward the end of the respective arm, one nucleotide at a time (Figure 3a).

Figure 3b shows that models built on longer regions of the 3′ arm have a better predictive power (60 nucleotides OOB=0.272) than models built on shorter regions of the 3′ arm (5 nucleotides OOB=0.608). We observed the opposite trend for the 5′ arm, where model performance worsens with length and the OOB error increases from 0.55 at 5 bases to 0.792 at 60 bases (Figure 3b). This indicates that the performance improvement is not due to giving additional degrees of freedom to the machine learning model. Adding to the already-established influence of ssODN length (29,30), we showed that the nucleotide composition of the 3′ arm influences HDR efficiency with symmetric ssODNs.

### Web service for predicting HDR efficiency

Combining information about the DSB frequency (guide) with information about HDR influencers (ssODN), we built a final model incorporating guide and 3′ ssODN information (M1), see Table 4. This model indeed identifies the largest number of high-efficiency HDR targets across our cross-validation folds (25 vs 23 (G1) and 22 (O3)). It hence has the highest recall rate although we observed the same OOB error compared to G1 (0.25). We use this model in our online webservice, CUNE.

**Table 4.** Metrics from the mixed model. This random forest model includes the 5’ oligo arm (O3) and the guide (G1). CUNE implements M1 to predict HDR efficiency of CRISPR-Cas targets.

| Model | Region | OOB error | ROC | Precision | Recall | Correct |
| --- | --- | --- | --- | --- | --- | --- |
| M1 | 3' Oligo & Guide | 0.250 | 0.91 | 0.883 | 0.8 | 25/30 |

CUNE identifies the optimal way to insert a specific point mutation at a genomic locus. With the efficiency of base editing on average higher than HDR, the service identifies which, if any, base editing system is applicable, using pre-established rules (5,11,31–33). CUNE also predicts the HDR efficiency for gRNAs and ssODNs around the specified locus.

The base editing component of CUNE is equivalent to BE-Designer (34), in that both tools identify guides that allow editing of a target using base editing. One difference is that CUNE allows the user to specify the target locus by genomic region, whereas BE-Designer requires the user to enter a target nucleotide sequence. Secondly, while BE-Designer covers a range of PAMs, CUNE specializes in the canonical SpCas9 PAM (NGG), as the most efficient SpCas9 PAM (35,36). Benchling is another service that provides support for identifying base editing targets (37). Like BE-Designer, Benchling allows the user to search within a region. Rather than scoring targets based on whether the mutation lies within the editing window, Benchling returns an efficiency score. However, Benchling only supports the classical base editor from Komor *et al*. (2016) and not more-recent base editors like BE3 (32).

In case base editing is not possible, our webpage provides the user with recommendations for high efficiency target sites for inserting point mutations with HDR. These recommendations are ranked by their predicted efficiency, according to our random forest model.

### Validation

Due to the traditionally low efficiency of HDR-mediated nucleotide edits compared to base editing, it has rarely been used in the literature so far, rendering us unable to find an independent dataset for validation. We hence created a second set of HDR targets from our more-recent mouse experiments. These thirteen targets, with nine as high efficiency and four as low, are distinct from our original targets. We used model M1 to classify these targets and compare the predictions to the truth labels. We observed three out of four low efficiency targets and eight out of nine high-efficiency to be classified correctly (Table 5). This quantifies the accuracy as 0.846 on unseen data for CUNE, as the first tool to computationally guide HDR target site selection.

**Table 5.**
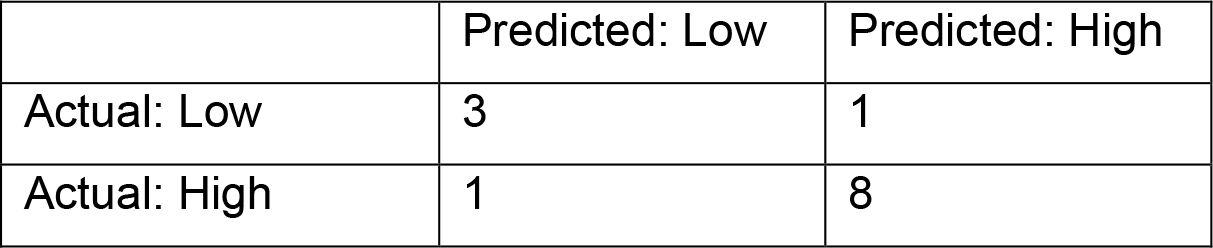
confusion matrix of the validation results on the thirteen unique targets. These targets were classified using the M1 model, with 11 out of 13 being predicted correctly. The prediction accuracy is 0.846.

|  | Predicted: Low | Predicted: High |
| --- | --- | --- |
| Actual: Low | 3 | 1 |
| Actual: High | 1 | 8 |

To quantify the theoretical improvement of choosing HDR targets using CUNE versus naively picking sites, we re-scored the targets in our dataset using CUNE. We compared the average efficiency of HDR targets predicted to be high-efficiency to the average of all targets in our dataset. This results in an 83% improvement in HDR efficiency for targets chosen using CUNE versus targets chosen randomly (predicted average efficiency of 0.528 compared to 0.288).

## Discussion

We set out to understand the factors that govern HDR-mediated point mutations. We aimed to create a computational tool that makes efficiency-improving recommendations for variables that are easy for the researcher to vary, such as ssODN design and guide. This is especially relevant, as the currently known factors to govern efficiency, such as cell type and locus (28), are usually fixed parameters for an experiment.

Using machine learning, we trained models on different features to investigate how such features influence the HDR efficiency. We chose to use the random forest algorithm as it enables us to quantify the contribution of each input feature (variable importance), as well as allowing us to model feature interactions, which provides insights into biophysical mechanisms. Random forests are also resilient to overfitting (38), which was crucial for some of our trainings scenarios containing more features than samples (e.g. G5).

We hypothesized that ssODN nucleotide composition would be an influencing factor on HDR efficiency, due to Watson-Crick base pairing between the ssODN and the DNA target being essential for inducing HDR-mediated point mutations. While the nucleotide content of the ssODN (O1) 5′ arm was unimportant, the content of the 3′ arm proved to be a major contributor to prediction accuracy.

The importance of the 3′ region is in agreement with the biophysical action of HDR. For a cell to proceed with HDR, the 5′ strands at the DSB are degraded (39). This process, known as 5′ → 3′ resection, results in 3′ overhangs at the DSB (Figure 4). Therefore, the 3′ region of the ssODN, being complimentary with one of the newly-formed 3′ overhangs, is the first region of the ssODN to interact with and bind to the target. We propose that if this occurs, HDR will continue regardless of the 5′ sequence, resulting in the poor performance of our 5′ ssODN models (Figure 3).

**Figure 4.**
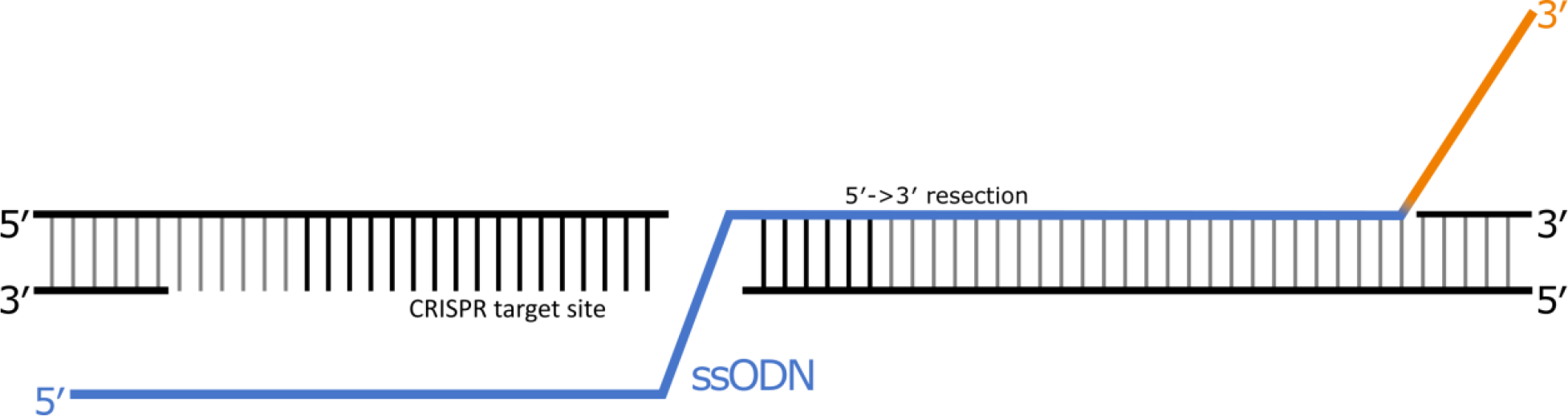
An ssODN (blue/orange) annealed to 5′-3′ resected DNA (PAM strand). ssODNs with regions extending beyond the resected DNA may require further processing or strand-invasion of the DNA target. The sequence composition of this region (orange) has a strong impact on HDR-efficiency.

Liang *et al*. (30) observed the optimal length for a 3′ arm to be 30-35 bases, which they base on the 5′ → 3′ resection at the DNA target typically creating overhangs of 30-40 bases. They suggest that arms extending beyond this region are accommodated by further target resecting, 3′ ssODN trimming or strand invasion of the target, while shorter arms can anneal directly to the target (Figure 4). We hypothesise the process that allows for longer ssODN arms is a highly sequence-specific, which we can investigate as our ssODN arms extend beyond the resected region. In support of the optimal length, we observed our prediction accuracy to temporarily plateau at 20 bases, before continuing to improve at 45 bases, all the way to 60 bases (Figure 3b orange). This indicates HDR-efficiency is especially sensitive to the nucleotide composition of the region beyond the resected DNA (Figure 4).

We hypothesised that the distance from the mutation to the PAM would contribute to our model’s accuracy, but this proved not to be the case in our training dataset. While we observed the expected inverse correlation between distance and HDR efficiency, reported in previous literature (27,30,40), the contribution to the computational model was low. It is likely that this is because of the unbalanced nature of this feature in our dataset. For example, Liang *et al*. observed HDR rates of below 5% at distances above 8 bases and rates of 10-30% at and below 8 bases. If we take 8 bases as the high/low threshold, a balanced dataset would require 50% of the samples to be up to (and including) 8 bases away, with 50% above 8 bases away. However, only 6 out of 30 (20%) of our samples are above 8 bases, limiting its impact in our model. This is a form of bias, as we selected gRNA targets primarily based on their close proximity to the PAM in this training dataset. Secondly, especially at short distances, the correlation between the distance and HDR can be quite variable. For the closest half of our targets, we see an average efficiency of 0.33. However, the interquartile range for these values is 0.478, nearly half the range of our targets (0 to 1).

Our work has resulted in the first computational method for designing efficient experiments for inducing point mutations using base editing and HDR (supp. figure 1). We have provided this as a web service, which will design ssODNs to induce user-specified point mutations. In addition, our webservice will also identify base editing targets using pre-existing rules.

## Methods and Materials

### Experiments

#### Ethical statement

All experiments were approved from the Animal Ethics Committee from the Australian National University according to code of practice of the National Health and Medical Research Council (NHMRC) in Australia (AEEC 2014/58 and A2017/44).

#### SgRNA single stranded oligonucleotide design and cloning

Mouse sequences (GRCm38/mm10 scaffold) were obtained from Ensembl (ensembl.org) or UCSC genome browser (genome.ucsc.edu). SgRNAs were designed close to the desired point mutation to replace with CRISPR-HDR and verified for off target effects using available online tools such as CRISPOR (19) or CCTop (20). SgRNA were designed as a gBlock from

IDT (Integrated DNA Technologies, Coralville, IA) encoding a U6 promoter, the SgRNA scaffold (crRNA and tracrRNA) with the following sequence adapted from Mali *et al*. (41).

5′-

TGTACAAAAAAGCAGGCTTTAAAGGAACCAATTCAGTCGACTGGATCCGGTACCAAGGTCGGGCAGGAAGAGGGCCTATTTCCCATGATTCCTTCATATTTGCATATACGATACAAGGCTGTTAGAGAGATAATTAGAATTAATTTGACTGTAAACACAAAGATATTAGTACAAAATACGTGACGTAGAAAGTAATAATTTCTTGGGTAGTTTGCAGTTTTAAAATTATGTTTTAAAATGGACTATCATATGCTTACCGTAACTTGAAAGTATTTCGATTTCTTGGCTTTATATATCTTGTGGAAAGGACGAAACACCGNNNNNNNNNNNNNNNNNNNGTTTTAGAGCTAGAAATAGCAAGTTAAAATAAGGCTAGTCCGTTATCAACTTGAAAAAGTGGCACCGAGTCGGTGCTTTTTTTCTAGACCCAGCTTTCTTGTACAAAGTTGGCATTA–3′

The 450 bp gBlock were TA cloned into a pCR2.1TOPO vector (Thermofisher Scientific, Waltham, MA, USA) according to the manufacturer’s instructions. SgRNAs were also chemically synthesised from IDT (Integrated DNA Technologies, Coralville, IA) as a crRNA and tracrRNA and assembled in a ribonucleoprotein complex with Cas9 enzyme according to the manufacturer’s instructions. 140 bp ultramer long Oligonucleotides from IDT at a concentration of 4 nmole were designed to target mutation of interest with 70 bp symmetric homology arms. Cas9 recombinant protein was purchased from PNA Bio (Newberry Park, CA, USA).

#### Mouse zygotes microinjection

C57BL/6Ncrl and Swiss Webster CFW/crl mice obtained from Charles River Laboratories and were maintained under specific pathogens free condition under 12/12 hrs light cycle and food and water were provided ad libitum. Three to five weeks old C57BL6/Ncrl females were superovulated by 5UI intraperitoneal injection of Pregnant Mare Serum Gonadotrophin (Sigma Aldrich, St-Louis, MI, USA) followed 48 hours later by 5UI intraperitoneal injection of Human Chorionic Gonadotrophin hormone (Sigma Aldrich, St-Louis, MI, USA). Superovulated females were mated with 20 weeks old stud C57BL/6N males. For microinjection, 45 to 48 hours following the second hormone injection, zygotes were collected from the oviduct. Pronuclear injections were performed under a DMi8 (Leica, Wetzlar, Germany) inverted microscope apparatus associated with micromanipulators and a Eppendorf femtoJet microinjection apparatus (Eppendorf, Hamburg, Germany). 50 ng/µl of Cas9 protein was complexed with 3 µM of crRNA and the tracrRNA or co-injected with 5 ng/µl of sgRNA plasmid and then mixed with the 50 or 100 ng/µl of single stranded Oligonucleotides and suspended in RNase and DNase free highly pure water (Thermo Fisher Scientific, Waltham, MA, USA) prior to the microinjection Microinjected zygotes were either surgically transferred into the ampulla of CFW/crl pseudo-pregnant females or cultured overnight at 37ºC in a 5% CO2 incubator and then surgically transferred at 2-cells stage of development.

#### Genotyping

DNA extraction was performed on ear punch from mouse pups over 15 days old mice using a crude DNA extraction. Briefly the ear punches were lysed in Tris-EDTA-Tween lysis buffer (50mMTris HCl, pH8.0, 0.125mM ethylenediaminetetraacetic acid (EDTA), 2% Tween 20) in addition to 1μl of proteinase K (20 mg/ml in 10mM Tris chlorate, 0.1 mM EDTA pH 8.0) and incubated at 56ºC for an hour before the DNA is being denatured at 95ºC for 10 minutes. Primers were designed to amplify the regions encompassing the sgDNA PCR were performed using Taq polymerase under standard PCR conditions. The PCR products were then purified with ExoSAP-IT1 (Thermo Fisher Scientific, Waltham, MA, USA) according to the manufacturer’s instructions. Sanger sequencing was performed at the ANU Biomolecular Resource Facility.

### Data collation

For the result of each CRISPR experiment, we manually collate the data into a spreadsheet. This results in a file with a row for each experiment, where each column stores a variable. We capture the following information:

- Guide sequence
- ssODN sequence
- Observed mutation (arbitrary or point)
- Distance of mutation from PAM

### Preparing the dataset

Where many CRISPR targets had one attempt to insert a point mutation, some targets had multiple attempts. This leaves us with samples sharing the same sequence as other samples. Therefore, to reduce any “sampling bias”, we merge duplicate targets. This involves summing up the number of attempts, arbitrary mutations, and point mutations. This leaves us with 57 rows of data.

We calculate the label by dividing the number of times we observed a point mutation for each experiment by the number of times we observed any mutation. Therefore, we discard any samples where we observed no mutation, leaving us with 30 rows of data. This results in a value from 0 to 1 for each experiment where 1 indicates 100% HDR, and 0 indicates 0% HDR (or 100% NHEJ). For binary classification (high or low HDR), we set a threshold. To balance the classes (highs and lows), we set the threshold to the median (0.199).

We use Python 3.5 with various packages including scikit-learn, Pandas, Numpy and Scipy. Using Pandas, we can read the data directly from the Excel file into a DataFrame. Here we encode values as integers for compatibility with the scikit-learn random forest library. We perform “one-hot encoding” on the discrete non-binary variables. That is, for each unique value in a column, we create new columns where the value is either 1 or 0. For example, the column that stores the first nucleotide from the PAM, “N_1”, is transformed to four columns four columns, “N_1_A”, “N_1_T”, “N_1_C”, “N_1_G”, where each column represents a specific nucleotide at that position.

### Statistical analysis

The primary metric we use is the out of bag (OOB) error. This metric takes advantage of one of the properties of the random forest algorithm, bootstrap aggregating (bagging). With bagging, each tree is trained on only a subset of samples. Therefore, each tree can be tested on the unseen samples to that tree. This is repeated for every tree throughout the training process. Finally, the average of the errors for each tree results in the OOB.

For further robustness, we partition our dataset using cross-validation (42) to evaluate our models. Using 5-fold cross validation, we split our dataset into five “folds”. We then train a model on four out of five folds, and test it on the fifth, repeated five times. This allows us to evaluate the prediction error with better generalization to novel data than a train/test set. We use “StratifiedKFold” (43) to create the folds, as it preserves the distribution of positive and negative samples.

We visualized the performance of each fold using receiver operating characteristic (ROC) curves (supp. figures 2-6). These plot the true positive rate against the false positive rate. The average area under the ROC curve for each model is presented in the corresponding tables.

### Scoring the model

The OOB error is the average of error values across each tree in the random forest. Because each tree is built using bootstrap aggregating (bagging), different sets of samples remain unseen to each tree. Therefore, the prediction error for each tree can be calculated on the unseen samples to that tree. The mean of these errors is the OOB error, where 0 represents perfect predictions and 1 represents random chance.

With a lack of HDR data in the literature, we validated the model on a more-recent data, completed after training our model. This data includes thirteen samples, being generated in the same way as our training samples (from experiments aiming to induce point mutations in mice). However, these new samples are the result of experiments targeting different genomic loci to those from our original data. We curated these samples in the same way as our original data to generate a dataset of features (guide and oligo nucleotide compositions) and truth labels (high or low efficiency). We scored this data using our mixed model (M1) and compared the predictions to the truth labels.

Our source code is available in supplementary item 2 and at: https://github.com/BauerLab/GT-scan2-Notebooks.

## Acknowledgments

We would like to thank Dr Jenna Lowe, Nikki Ross, Jing Gao, Nay-Chi Khin and Lora Starrs for their technical assistance. This work was supported by the National Collaborative Research Infrastructure (NCRIS) via the Australian Phenomics Network (APN). We thank Alex Whan and Daniel Layton for critical reading of the document.

